# Exovelo: Cell States Velocity Estimation Without Parameter Fitting

**DOI:** 10.1101/2025.07.02.662720

**Authors:** Yiftach Josef Kolb, Laleh Haghverdi

## Abstract

Reliable estimation of cell state velocities (in gene expression space) has been largely hindered by model presumptions which may not be true for all genes, followed by a highly inaccurate procedure of inference of the assumed model parameters and visualization artifacts. Here we introduce Exovelo, a velocity estimation method without the conventional gene-wise parameter fitting step that actually allows de novo learning about the single-cell dynamics of cell differentiation beyond geometrically inferred pseudotime methods. By adapting an appropriate data rescaling, Exovelo makes the distributions of current and future cell state estimates to overlap, then visualizes the expected cell state displacement in a joint embedding. Exovelo estimates velocities from pairs of single-cell unspliced-spliced or metabolically unlabeled-labeled mRNA data, as we demonstrate on simulation as well as datasets form cell-cycle and hematopiesis.

## 2. Introduction

In the universe of single cell RNA sequencing (scRNAseq) every cell can only be sequenced once and it is destroyed in the process. Consequently an individual cell state’s velocity cannot be directly observed. However current sequencing techniques can distinguish between different species of transcripts (spliced vs unspliced, or labeled vs unlabeled) and that distinction conveys temporal information about the RNA transcript’s unique molecular identifier (UMI). The first single-cell rna velocity estimation method was introduced by [1], with several follow-up methods as well as critical views on the existing methodology [2, 3].

In this paper *cell state* refers to, depending on context, either the vector raw of RNA transcripts of a cell, or to some transformation of it which may include some or all of the following stages: gene selection, normalization, rescaling and log–transformation, PCA or some other linear dimensional reduction. *Cell state velocity* or *RNA velocity* is the (estimated) vector with direction and magnitude of change of a cell state, or the displacement in a cell state over a short time interval.

Current scRNAseq technologies in conjunction with tools such as Velocyto [1] allow the calling of each UMI as (likely) spliced or unspliced. Unspliced RNA, also referred as precursor mRNA, is converted to spliced RNA which is the final, mature form of the transcript by the process of splicing. Under the assumption that splicing half life is significantly shorter than the mature RNA half life, an unspliced RNA transcript is very likely to be newer than any spliced transcript.

The second type of quantification is made possible by metabolic labeling [4] in conjunction with tools such as Dynast [5]. Without delving into the technical details of this techniques, the end result is that each UMI in the dataset is either ‘labeled’ or ‘unlabeled’ where labeled implies that the molecule had been transcribed within a known time interval before sequencing (the labeling time interval), and an unlabeled molecule must had been transcribed prior to labeling start. Labeled molecules are also called ‘new RNA’ and unlabeled ones are called ‘old RNA’.

Since the inception of the RNA velocity field by [1] a number of inference methods and their associated tools have arisen. Most notable examples to our knowledge are Velocyto [1], scVelo [6], VeloVAE [7], Dynamo [8] and *κ*-velo / eco-velo [2]. With the exception eco-velo, all the other models have an explicit underlying assumption that the dynamics of spliced and unspliced RNA (and for the tools which also infer with labeled data, old and new RNA), follow a system of ordinary differential equations (ODE) with fixed kinetic rate parameters. Some of these models may add additional constraints such as on the number of switches a gene can undergo in a dataset. In “Eco-velo” there is no parameter fitting, however it has an underlying assumption that degradation rate *γ* equals splicing rate *β* and under this assumption unspliced RNA is proportional to spliced RNA from 1*/γ* time units in the past. Moreover, Ecovelo similar to the previous methods, treats velocities as “arrows” rather than vectors with defined direction as well as magnitude.

Here we propose “Exovelo” which advances the parameter-free inference concept to a mature tool ready for practical use. In the metabolic labeling case Exovelo normalized and scales the total *R* and old *O* RNA so that after proper transformation 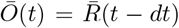 where 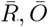, are the transformed, scaled modalities, *t* is the unknown latent time, and *dt* is the labeling time. Similarly in the spliced/unspliced case Exovelo transforms and scales the spliced *S* and total *R* modalities so that 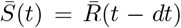. In this case *dt* is related to splicing half life but is not exactly known.

For visualizing the inferred velocity (more correctly displacement over an implicit or explicit time interval) Exovelo creates joint embedding (JE) which is in our view the most faithful visualization method, but also can create neighbor based which in our experiments produces qualitatively similar results to JE (figure 1 a-d).

**Figure 1.**
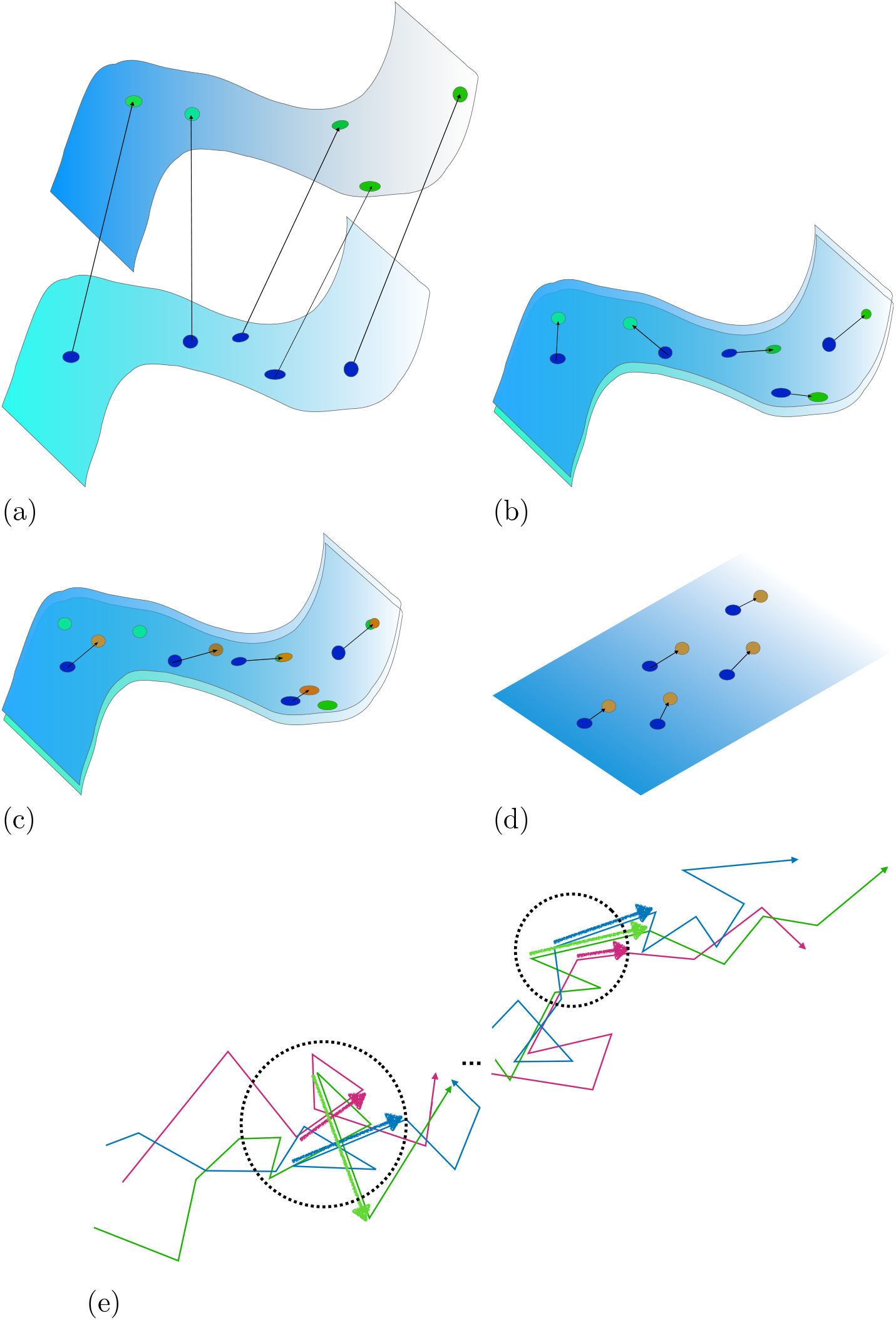
Schematic summary of Exovelo methodology. (a) Bimodal high dimensional unscaled data. (b) Rescaled high dimensional data. (c) Mean field velocities based on neighbor estimation (see also (e)). (d) Reduced dimension visualization by joint embedding. (e) Assuming that each cell and its estimated displacement is sampled from one single trajectory, mean of displacement vectors of a cell’s neighbors approximates its drift component, and their variance approximates its diffusion magnitude.

If we assume a diffusion-drift model for cell differentiation as proposed for example in [9], Exovelo can optionally utilize that assumption to infer the drift component. To that end it uses a cell’s neighbors velocities as a sample set for possible velocities the cell could have in order to infer a mean field velocity for each cell representative of its drift component. Similarly the diffusion magnitude can be estimated by calculating the variance of neighbouring cells velocities (figure 1e).

In the following, we tested Exovelo on simulated data where ground truth is known and confirmed it infers correct velocities even in a case where it cannot be done with kinetic parameter inference using scVelo. Our test on the human retinal pigment epithelial-1 (RPE-1) dataset from Battich et. al [10] shows that Exovelo produces very similar, and biologically correct results when used on spliced/total and on unlabeled/total modalities, supporting the feasibility of Exovelo on datasets which only have the splicing information and are not metabolically labeled. Finally we tested Exovelo on bone marrow dataset [11] which is more challenging. We argue that even in this case and considering the asymmetric differentiation and multiple possible differentiation paths of hematopoietic stem cells Exovelo infers biologically relevant cell state velocities.

## 3. Results

### 3.1. Simulated trajectory dataset

We tested Exovelo on simulated data where ground truth of time and splicing modalities are known. The data comprises from a set of trajectories which all share the basic transcription state at each time point but differ by random diffusion and random variations in the rates of transcription degradation and splicing. In figure 2a–b we illustrate how per gene rescaling of the two modalities can capture the shift between the current and future state of the cell both in upregulation and downregulation phases of gene expression. Using the multi-dimensional (i.e., several genes) displacement vector, Exovelo is able to infer the correct velocity direction (figures 2c and 2d). The mean Pearson correlation coefficient between the real displacement and inferred displacement is 0.83 for Exovelo. We also tested scVelo on the simulated data (figure 2e). In our tests scVelo inferred the correct trajectory direction although with tiny velocities. That was somewhat surprising given that the simulated transcription dynamics doesn’t match scVelo assumptions with respect to constant rate parameters. In simulations where there are relatively fast changes in gene dynamics as shown in the plots scVelo didn’t consistently infer the trajectory direction.

**Figure 2.**
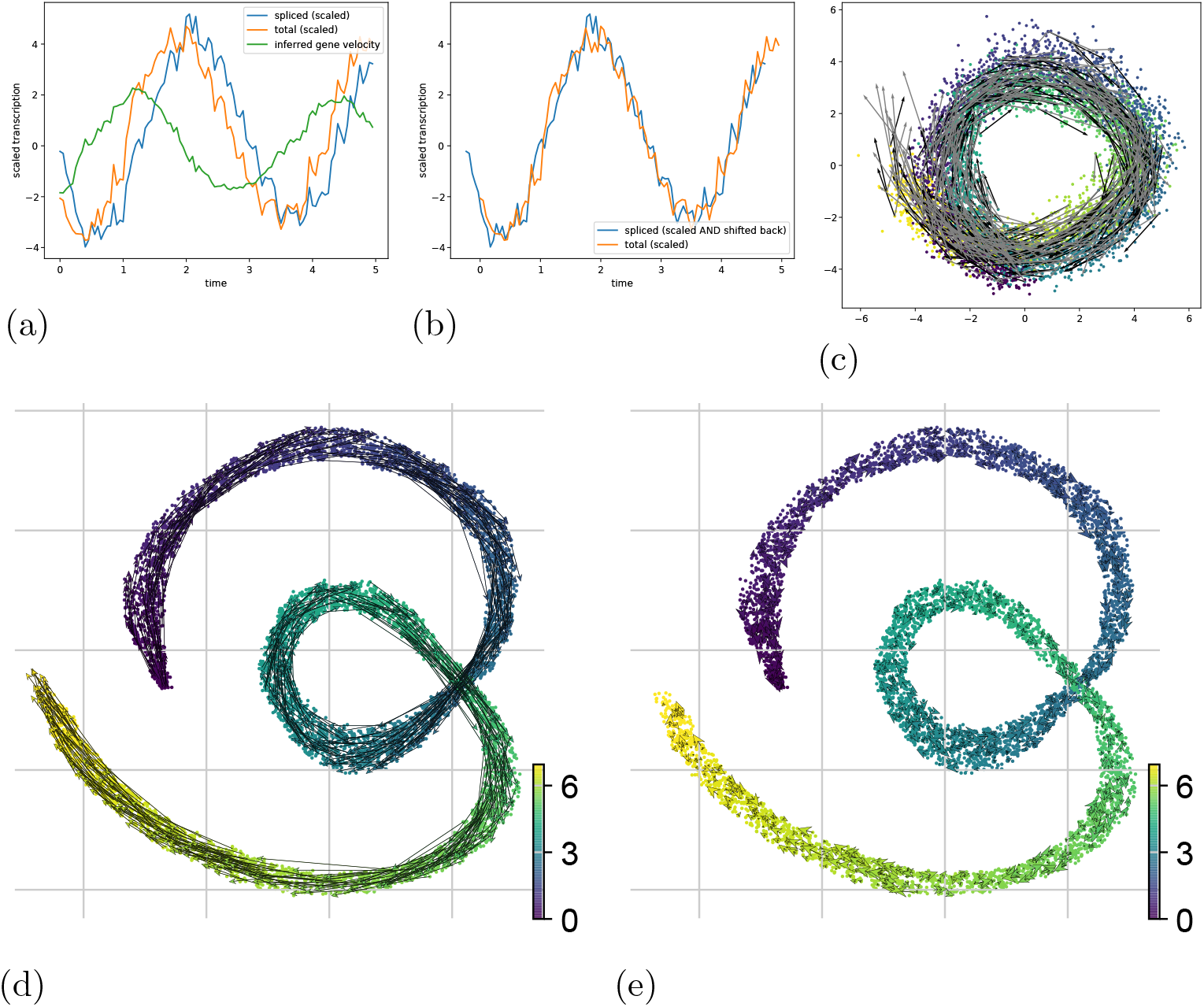
Simulated data of quasi–cyclical trajectory. (a) After proper scaling, total RNA reflects the ‘future’ of spliced RNA in both its up– and its down–regulated phases. (b) The spliced modality shifted leftwards by splicing half life (log(2)*/β*) very closely matches the ‘total’ modality when *β > γ*, (*γ* is the RNA degradation rate). The assumption of constant rates is not a necessary requirement for Exovelo, however in case where they are the phase difference is proportional to the splicing half-life. (c) JE PCA velocity plot. gray arrows: actual displacement. black: Exovelo’s estimated displacement. The mean Pearson correlation coefficient between the two vector fields is about 0.83. (d) JE UMAP with inferred velocities by Exovelo shown in actual scale. (e) Same JE UMAP with velocities inferred and projected by scVelo. In all the quiver plots cell time is indicated by viridis color map (purple: early, yellow: late).

### 3.2. RPE1 dataset

In the case of metabolically labeled data old (unlabeled) RNA is literally the remaining total RNA from time point (*t − dt*) where *t, dt* are respectively the sequencing time and the labeling duration. Exovelo applies the same method to spliced/unspliced modalities and it takes spliced RNA is a proxy for ‘old’ RNA. However *dt* is unknown in the splicing case, and the time scale itself varies between genes and possibly even for the same gene between cells.

The dataset of immortalized retinal pigment (RPE1) cells from Battich et al. [12] contains information both on splicing modalities as well as metabolic labeling with 6 labeling times. This is a ‘cell cycle’ dataset as the RPE1 cells proliferate but don’t differentiate.

To demonstrate the importance of gene-wise rescaling in Exovelo, we order the cells according to their relative cell cycle position (using the values from the original publication), and for two example genes. It can be seen in Figure 3 (for TPX2 gene), and also in supplementary figure A1 (for TOP2A gene), that for both unlabled-labled and unspliced-spliced datasets, after gene-wise rescaling of the old and total mRNA counts the two modalities become comparable. The difference between total RNA (the ‘future’ cell state) and Old RNA when the gene is down regulated tends to be negative as it should be in this case, and it tends to be positive when the gene is up-regulated. In the splicing case the phase shift between spliced (‘past’ cell state) and total RNA (the ‘future’ state) is much smaller but still exists. Moreover rescaling the two modalities so that they become comparable is in some sense the best we can do. The phase shift ‘should be there’ after rescaling however technical issues such as inaccurate spliced/unspliced calling or very low unspliced count can make the phase shift disappear in some regions. Note that the ordering of cells in Figures 3 and A1 are only for demonstration of rescaling effect such that the difference (total RNA - old RNA) or (total - spliced RNA) is in sync with the gene phase (negative when down regulated, positive when up-regulated). Exovelo itself neither uses any ordering of the cells nor any kind of sliding window over the cells.

**Figure 3.**
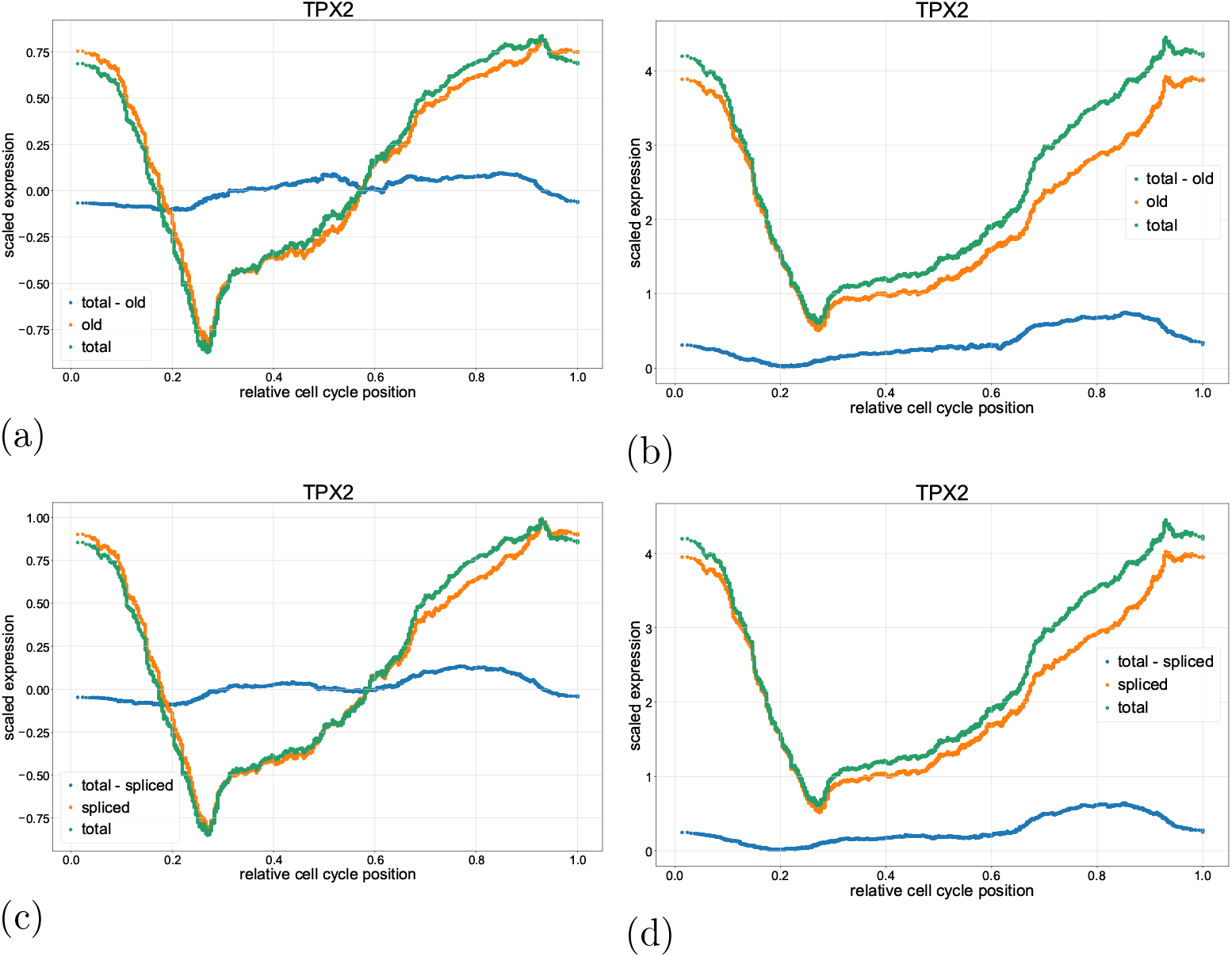
TPX2 gene expression over estimated cell cycle position. Labeling modalities: (a) after — and (b) before, gene-wise rescaling. Splicing modalities: (c) after and (d) before, gene-wise rescaling. The total-old curve (blue), represent the gene’s velocity. During the gene’s downregulation phase, the total mRNA amount (i.e., new) follows the old (negative gene velocity), whereas during the upregulation phase, the total surpasses the old (positive gene velocity). This pattern can be correctly seen only after gene-wise rescaling.

The velocity analysis results are shown in figure 4. scVelo [6] was used for plotting of displacements inferred by Exovelo. The results are consistent between labeling or splicing information. The cells appear to be cycling except for a subset of cells in the G1 phase which seems to be exiting the cell cycle (perhaps to a resting g0 phase). The embedding in figures 4a and 4b were produced by joint embedding on of the modalities which have been rescaled by Exovelo. Inferred displacement from splicing and from labeling information show high correlation and produce similar drift directions; figure 4c shows the correlation of their respective first principle compo-nents. As seen in figure 4d inferred displacements of cells with longer labeling duration tend to have larger magnitude than that of cells with shorter labeling time, as expected. The growth in magnitude with increasing labeling time is sub-linear however the data had been log-transformed.

**Figure 4.**
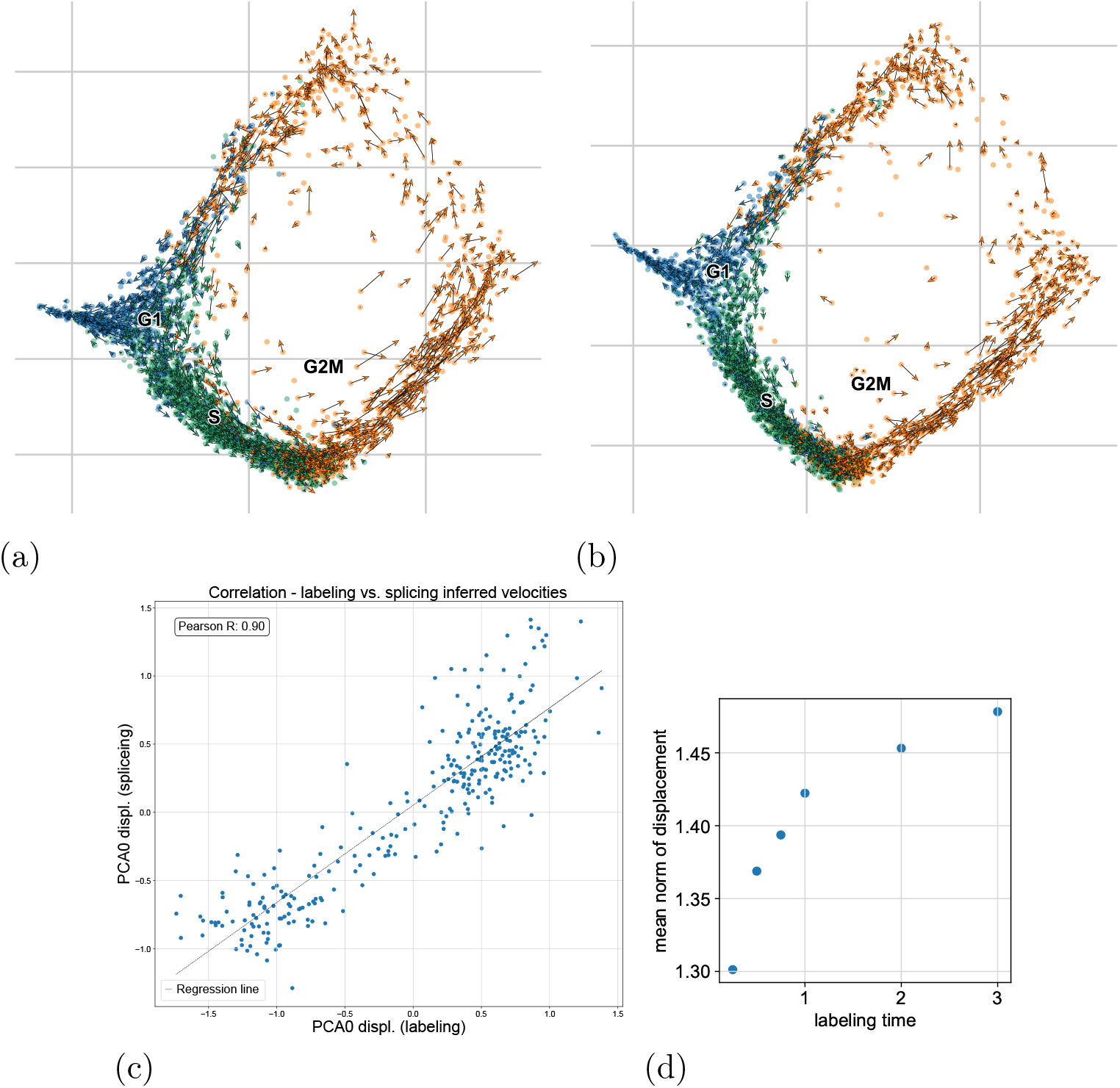
Cell state velocities of RPE1 cycling population. (a) JE diffusion map with velocity inferred from the labeling information. (b) JE diffusion map with velocity inferred from the splicing information. (c) Inferred velocities in the labeling vs. splicing case, shown on the first principle component with Pearson correlation coefficient of 0.9. (d) Vector norms of inferred cell state displacement tend to be larger with longer labeling duration.

### 3.3. Human bonemarrow dataset

We next tested Exovelo on hematopoietic stem cells and progenitors (HSCs) dataset [11] which has been analyzed also in [2]. Hematopoietic stem cells have proved challenging for RNA velocity tools [8, 15]. Moreover one needs to consider what is the actual ground truth velocity and whether it can be inferred at all in this case. HSCs are multipotent stem cells which give rise to all blood cell types. When a mother HSC stem cells divides, its daughter cells, initially very similar to each other, can each become any of the following cell types: 1) HSC. 2) multi potent progenitor cell (MPP). 3) common lymphoid progenitor (CLP) 4) common myeloid progenitor (CMP). Additionally there is a distinction between long term and short term HSCs with the latter being more committed to differentiation. We show schematically in figure 5a, such a scenario that cells from a similar state neighbourhoods may experience different drift forces towards multiple directions.

**Figure 5.**
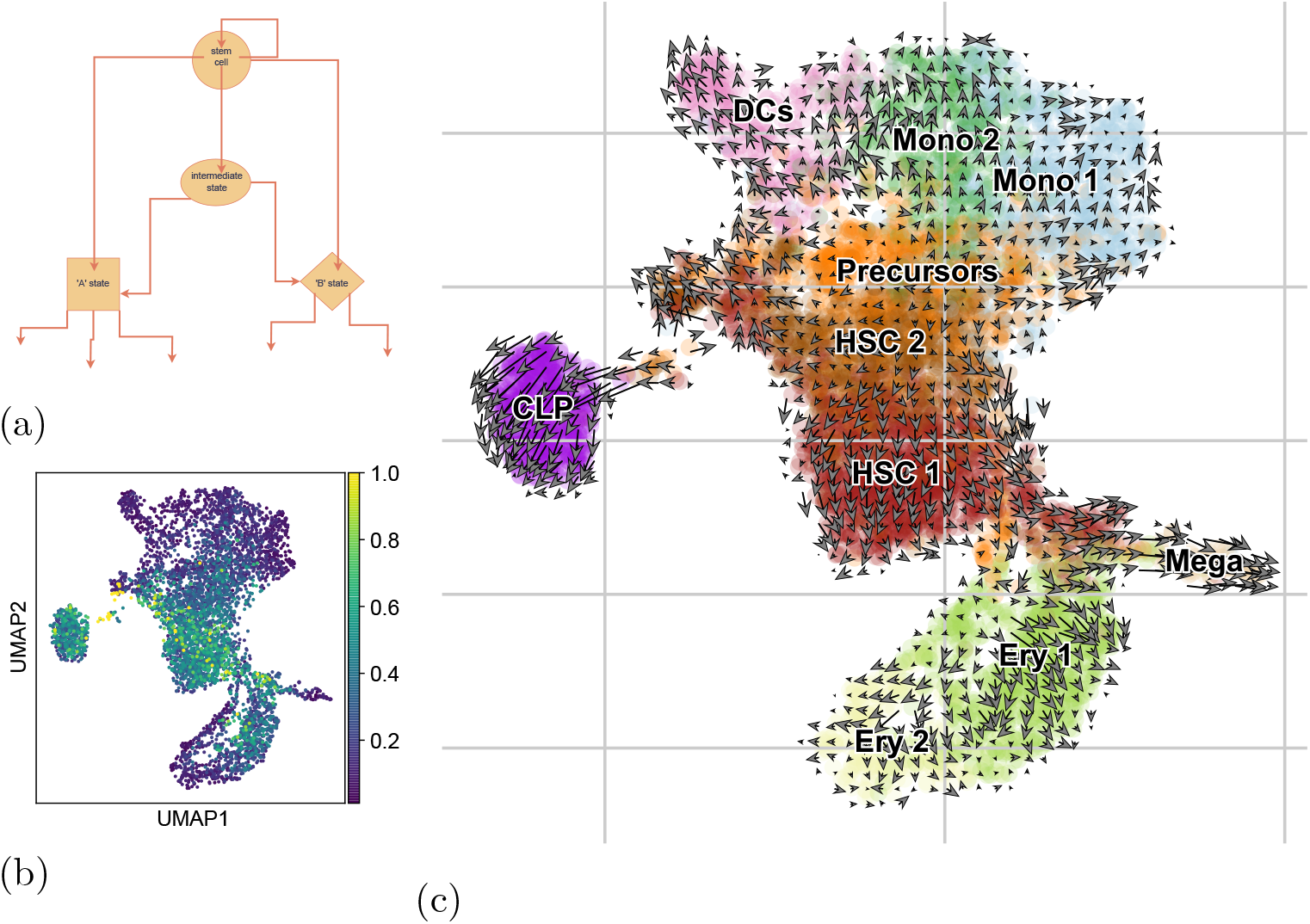
Exovelo application on Hematopoietic stem cells and progenitors from human bone marrow. (a) An illustration of cell differentiation process. A stem cell has the capacity to differentiate (by cell division) into any 2-combination of the states which it directly connects to. (b) Velocity variance is highest among the HSCs and tends to decline with differentiation. (c) Exovelo JE UMAP. HSCs and precursor cells seem to be going in all sort of directions including “in reverse” whereas more differentiated cells tend to go further down their branch.

HSCs are capable of self renewal. MPPs may lack self-renewal [16] but can further differentiate into either the myeloid or lymphoid paths. MPPs (resp CLPs) are committed to the myeloid (resp. lymphoid) path and give rise to any of the various cell types in each lineage.

Because of the self renewal capability of HSCs and the multiple possible differentiation paths the ‘true’ RNA velocity field need not always be directed downstream the path of differentiation but also in reverse (daughter cells returning into HSC stem cell state) or undetermined (velocities in many directions cancel each other out). Clusters of cells with multiple differentiation paths such as the ‘precursor’ cell cluster may have very different transcription velocities at the single cell velocity resolution, hence showing larger displacement variance than the more differentiated downstream populations (figure 5b and 5c). We tested Exovelo on another Bone marrow dataset [17] and observed similar patterns with larger diffusion near the pluripotent state and larger drift downstream (figure A2 in the supplementary figures section).

## 4. Methods

### 4.1. Main concepts of Exovelo

In this section unless otherwise stated capital letters represent matrices of shape (*n, m*) where rows are observations (cells) and columns are variables which depending on context are either genes or variables in a reduced dimensions representation such as PCA. Specifically *R, S, O* represent the total, unspliced, and old (unlabeled) RNA. Computations are done element-wise unless otherwise stated. Assume for simplicity that all cells are sequenced at the same time *t*. We consider the data matrices as functions of time, so that *R*(*t*) indicates the dataset at time *t*. There is only information for one time point *t* but there are 2 modalities either *R*(*t*), *O*(*t*) or and *R*(*t*), *S*(*t*).

### 4.2. Metabolic labeling case

Consider the metabolically labeled data case. Let *dt* be the labeling duration. Then *O*(*t*) represents all the remaining total RNA which had been produced prior to the labeling begin at time (*t − dt*).

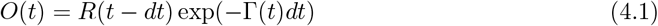

Where Γ(*t*) is a cell over gene matrix representing the RNA degradation rate for each cell for each gene at time interval [*t − dt, t*]. Had there not be any degradation we would have *O*(*t*) = *R*(*t − dt*). Exovelo seeks to transform *R*(*t*) and *O*(*t*) in a way that equation 4.1 will hold with Γ ≡ 1. If RNA degradation rate is the same constant for all cells and for each gene in the dataset the matrix Γ(*t*) becomes a (1, *m*) shaped constant vector *γ*. equation 4.1 becomes:

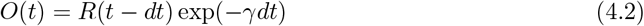

In equation 4.2 every gene in *O*(*t*) is proportional to the corresponding gene in *R*(*t − dt*) with fixed proportion *O*(*t, g*) ∝ *R*(*t − dt, g*). In particular, the mean and standard deviation of every gene in *O*(*t*) are proportional with the same fixed proportion to the corresponding gene’s mean and standard deviation. Let *Ō* (*t*) and let 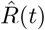 be the rescaling of *O*(*t*) (resp.) *R*(*t*) so that each gene is rescaled to 0 mean and 1 standard deviation. Then from the above discussion we have:

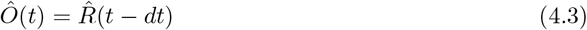

And from equation 4.3 we obtain the displacement estimation on the standardized data which is a proxy of the RNA velocity:

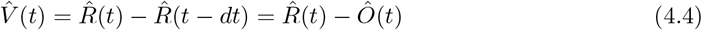

In case *O, R* are log transformed the proportional relation becomes a translation but after rescaling (which amounts to centering followed by division by the std) equations 4.3 and 4.4 still hold after log transformation followed by rescaling.

### 4.3. Splicing case

The splicing case is done exactly the same as the labeling case with *S*(*t*) used instead of *O*(*t*). In this case *dt* is unknown and in principle it is different for each gene however we experimentally observe that this approach works. Suppose that the splicing rate is greater than the degradation rate *β > γ* and in this case *dt* is in the order of magnitude of ≈ 1*/β*. Under these assumptions Any unspliced molecule from time *t − dt* or earlier is very likely to be spliced by time *t* and therefore *S*(*t*) consists substantially of all the remaining total RNA that existed at time *t − dt*:

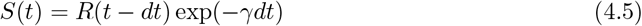

And from 4.5 the exact rescaling procedure as in the labeling case leads to:

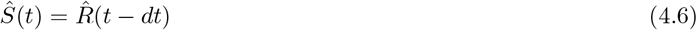

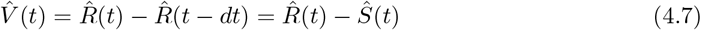

To see what is the optimal time scale *dt* = *τ* for constant rates *α, β, γ* the following rational is use: *τ* must optimize the following two conditions:

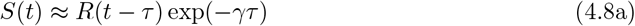

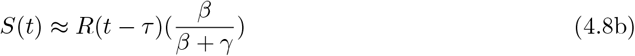

The first condition 4.8a is the same as 4.6 and the relation in the second condition 4.8b comes from the ratios of *S* to *R* at their steady states limits. It follows therefore that:

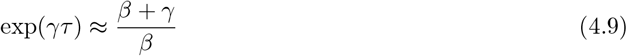

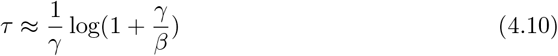

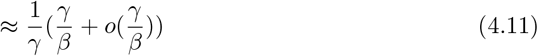

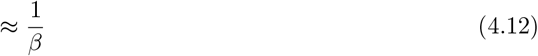

The calculation above used Taylors approximation for log(1 + *γ/β*) assuming that *γ/β* is small.

### 4.4. Data transformation and rescaling

Let *X* and *Y* be the raw data cell over gene matrices where *X* represents the past modality (either unlabeled or spliced) and *Y* the future modality (total RNA in either case). These undergo typical preprocessing— filtering cells and genes with low count, sub-setting for highly variable genes etc. and then size normalization. All these transformations preserve the proportionality of means between *X* and *Y*. Log transformation is a subsequent, optional transformation before rescaling. In metabolically labeled datasets which include more than one labeling time, rescaling is done separately for each labeling time.

### 4.5. Diffusion-drift dynamics

#### Fokker-Planck equation and stochastic differential equations

Here we follow the formulation from [18]. It has been suggested for example by [9] that the time development of the distribution of expressed RNA for a population of differentiating and proliferating cells P(***x***, *t*) can be described by the Fokker-Planck equation 4.13.

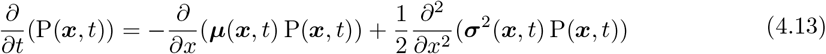

More generally equation 4.13 describes the motion of a *random process* ***X***(*t*) — that is— for every *t*, ***X***(*t*) is a random vector with distribution P(***x***, *t*). The stochastic movement of ***X*** can be described by the following stochastic differential equation:

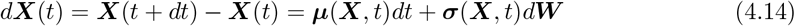

Where ***W*** is a *Wiener process*— a random process that models the integrated effect of noise or diffusion and without it equation 4.14 would be a deterministic ODE. ***W*** has by definition the following property:

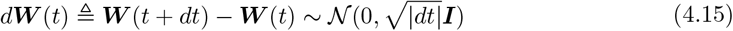

Where ***I*** is the identity matrix.

In the scope of this paper ***X*** represents the RNA expression of a random cell in the dataset (after feature selection, scaling etc.). In a scRNA-seq dataset there is no explicit information about the latent time *t* and had we tried to infer *t* from the data it would inevitably be a function of the cell state ***x***. Moreover it is reasonable to assume that cell state transition depends directly only on the current cell state and not on the latent time. The model for *d****X*** is therefore reduced into *t*-independent form:

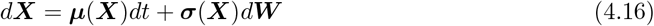

#### Estimating the drift component

Exovelo obtains an estimation for every cell states displacement which contains both the drift and the diffusion components. Specifically in the labeled case It uses 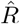, *Ō* and equation 4.4 as samples for ***X***(*t* + *dt*), ***X***(*t*), *d****X***(*t*), or 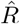, *Ŝ* and eq 4.7 in the splicing case.

In order to estimate the drift component Exovelo then applies equation 4.16. For every cell, the displacements of it *k* nearest neighbors are taken as a sample for *d****X***(*t*) and the estimated drift component *d****X***(*t*) is the mean of these displacements.

### 4.6. Visualization

#### Joint embedding

Since Exovelo infers two cell states (past, future) for each cell which are first class citizens the most reliable visualization for these states is a joint embedding (JE). In a JE an embedding is created from the 2*n* cell states (*n* cells × 2 cell states). The ‘velocity’ or more accurately the displacement is the vector from the embedded past state to the embedded future state of each cell. One advantage of a JE is that it is easier to catch failure, for example if Exovelo fails to properly rescale the two modalities then a strong batch effect will be noticeable in JE.

#### Projection

If projection of high cell state displacement into an existing embedding is desired we propose the following method. Let *X, Y* be rescaled cell × (high dimension) data matrices representing the ‘past’ and ‘future’ cell states of each cell. Let *p*: *X* → *Z* be any low dimensional projection of *X* e.g. a UMAP. An arbitrary cell *c* in the bimodal data can be represented as a vector pair (*x*,_*c*_, *y*_*c*_) of its past and future states. Let *N*_*k*_(*x*_*c*_, *X*) represent the *k* nearest neighbors in *X* of *x*_*c*_ ∈ *X* and *N*_*k*_(*y*_*c*_, *X*) the *k* nearest neighbors of *y*_*x*_ again in *X*. Since *y*_*c*_ is the ‘future’ of cell *c N*_*k*_(*y*_*c*_, *X*) is the set of its future neighbors. The projection we suggest is the **shift in the centers of masses** between the two sets, namely the low dimensional representation *v*(*x*_*c*_) of the velocity of *c* is:

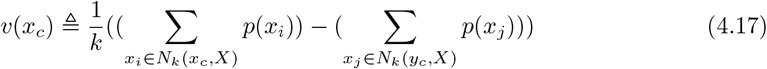

As is the case with any velocity visualization these can be smoothen by taking mean over the embedded velocities of a cell’s nearest neighbors. In our experiments this projection method produces similar velocity plot to JE when projected into the same embedding (supplementary figure A3).

## 5. Discussion

The introduction of single-cell RNA-velocity in [1] was widely praised by the computational biology community, as it would enable the study of dynamic processes at the single-cell level, surpassing the limitations of pseudotime methods, which only capture collective dynamics trends. However existing approaches assume model dynamics details that are non-essential for velocity estimation, but make the inference procedure non-transparent, error-prune and costly. Exovelo aims for estimation of cell’s expected displacement in a given time interval, hence turning the cell state velocity computation framework around, such that the dynamics can be meaningfully characterized from estimated velocities rather than vice versa. Thus, in subsequent research, models of cell differentiation dynamics (e.g., smooth or stochastic jump processes, asymmetric cell division, etc.) could build on such less-biased estimated velocities.

Cell state embedding and cell state velocity should be linked and in our opinion a joint embedding is the most faithful low dimensional representation for that reason. We caution not to rely on velocity projection into an arbitrary embedding which might be based on different feature selection, different normalization and metrics, and different local structure that was used for velocity inference. For example there could be a situation where a cells’ nearest neighbors as calculated used for projection are located in different clusters in different region of the embedding. On the other hand if the embedding has the same local neighborhood as the high dimensional manifold on which displacement is inferred then the Exovelo projection is very similar to a its joint embedding, at least in our tests and also logically the computations in both cases are very similar (supplementary figure A2)

There are some datasets that everything you try on them works. One such dataset is the embryonic endocrine progenitors and differentiated pancreas cells [19]. Another such dataset is of the human retinal pigment epithelial-1 [10]. The reason why these dataset are so “easy” may partially be technical, and partially that they are biologically less complicated than for example Hematopoiesis. The pancreas cells have a simple lineage tree with basically a single split from progenitors into the differentiated types. The RPE1 cells only cycle and don’t differentiate. On the other hand with hematopiesis the lineage is much more complicated. There are multiple branches, cells capable of self-renewal and possibly de-differentiation therefore the trajectories are not one-way and distinct.

In future work, Exovelo’s approach could potentially be improved upon by adapting a more sophisticated and fine-grained transformation, for example by factoring different cell clusters separately.

RNA velocity still faces challenges both on the inference method level as well as the data origination level. While this paper deals exclusively with the former challenge, it is not a given that datasets with splicing or labeling information are sufficiently accurate.

## 6. Code and data availability

The code including Exovelo python package and all three datasets mentioned in this paper are available in the [HaghverdiLab github repository](https://github.com/HaghverdiLab/exovelo_paper).

The RPE1 dataset [10] was downloaded using Dynamo [8] and can be [downloaded here](https://figshare.com/ndownloader/files/47439641).

The human bone marrow dataset [11] was downloaded using scVelo [6] and is [available here](https://ndownloader.figshare.com/files/27686835).

The raw second Haematopoietic dataset [17] is accessible in the Gene Expression Omnibus (GEO) under accession code GSE226824.

Exovelo is implemented in Python3.11 [20] and relies on Numpy [21] and Scipy [22]. Scanpy [13] is used for input/output and preprocessing of sequenceing data. Visualizatio relies on scVelo [6] Scanpy and Matplotlib [23]

## A. Supplementary

### Individual gene plots

**Figure A1:**
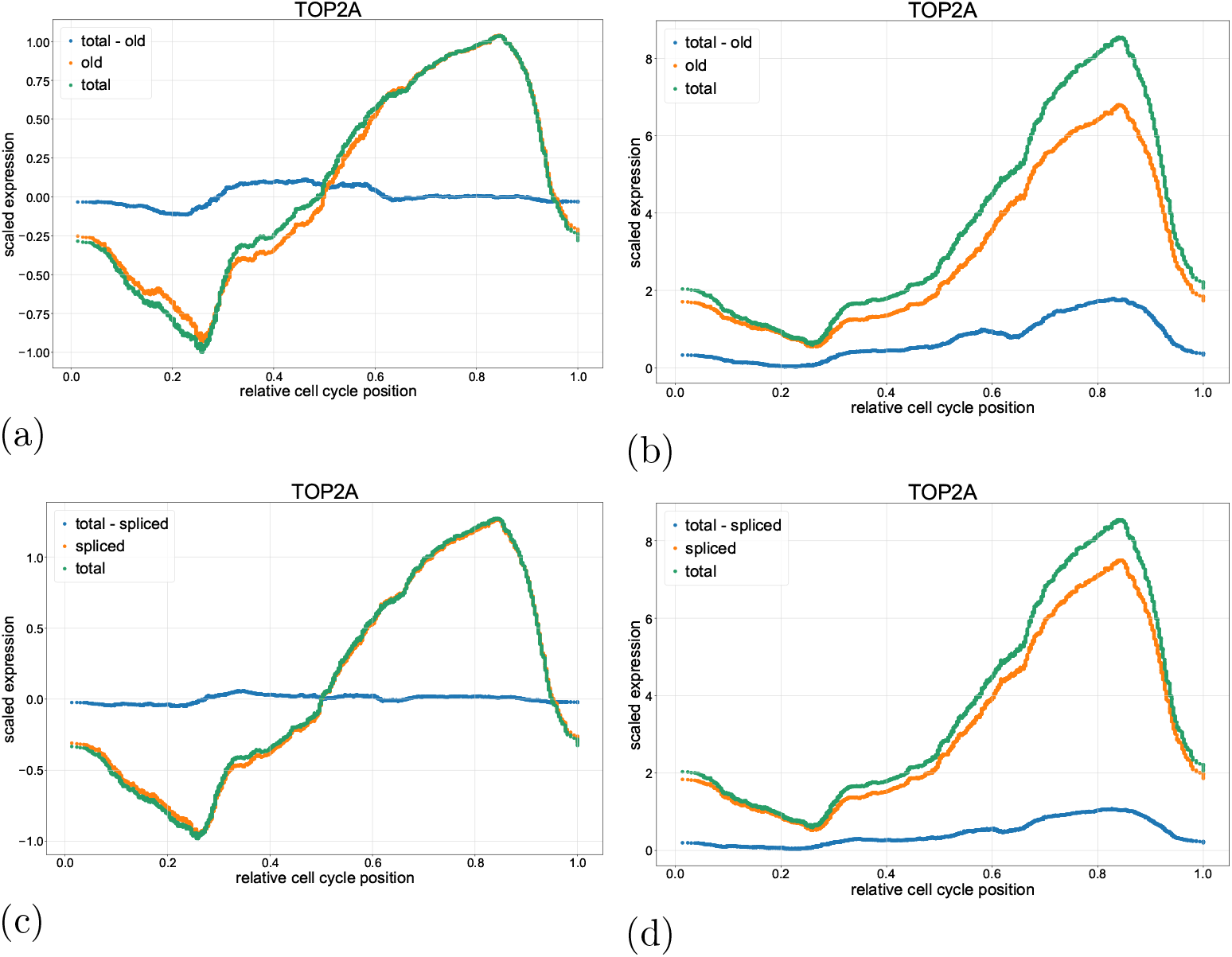
TOP2A gene expression over estimated cell cycle position. Labeling modalities: (a) after — and (b) before, gene-wise rescaling. Splicing modalities: (c) after — and (d) before, gene-wise rescaling.

### Additional bone marrow dataset

**Figure A2:**
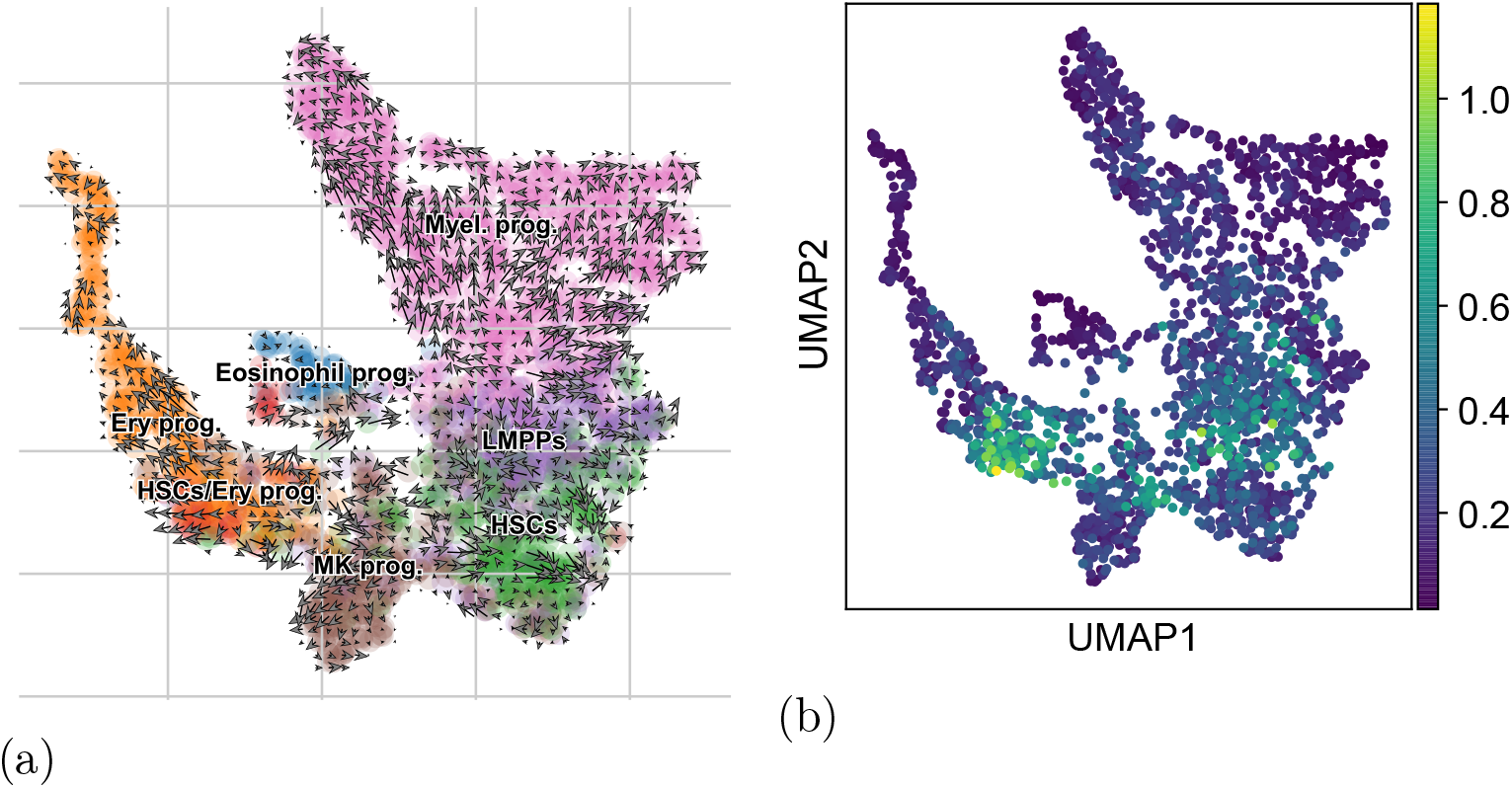
Exovelo application on HSPCs dataset of [17] (a) JE (smoothened) umap of Bonemarrow datasets from [17] showing mean field velocities. (b) shows the velocity variance of the same dataset. Similar to the first HSPCs data set [11] (see Figure 4 of the main text), here too velocity variance is greatest for the stem cells population (HSCs) and declines in conjunction with cells progress in differentiation. In both datasets there are semi differentiated precursor cells with near zero mean field velocity which have non-zero variance. In other words there are cells in these clusters which go in opposing directions and therefore no particular tendency is observed. Cells in the more differentiated parts seem to move down their respective branch.

### Comparison of projection of velocity vector onto the old states embedding with JE

**Figure A3:**
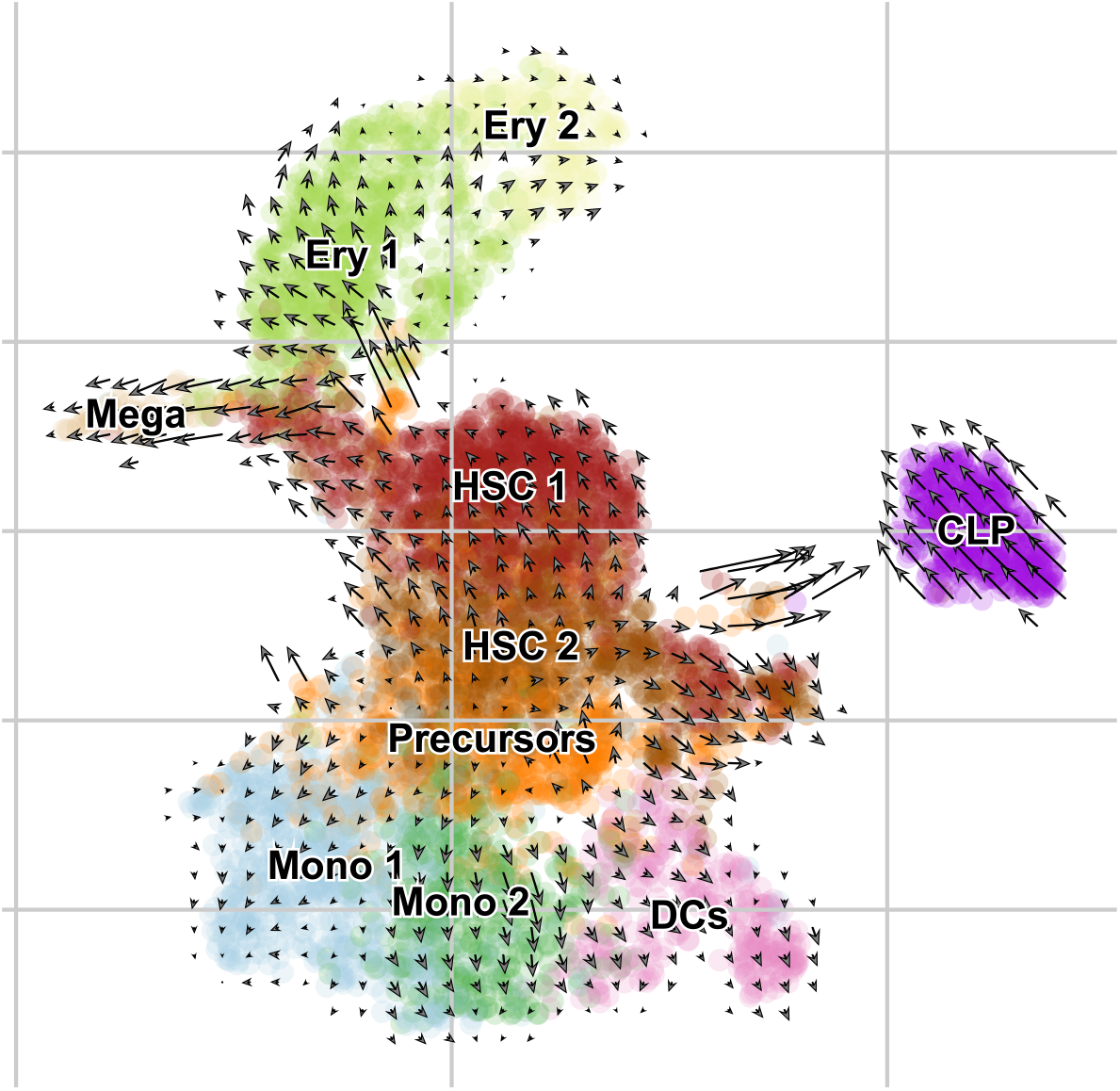
Exovelo projection of inferred displacement into a UMAP constructed just for the spliced modality. If the high dimensional inferred displacement is the same, Exovelo projection produces very similar result to Exovelo JE. Compare with figure 5c

